# Pipeasm: a tool for automated large chromosome-scale genome assembly and evaluation

**DOI:** 10.1101/2024.10.21.598381

**Authors:** Bruno Marques Silva, Fernanda de Jesus Trindade, Lucas Eduardo Costa Canesin, Giordano Souza, Alexandre Aleixo, Gisele Lopes Nunes, Renato Renison Moreira Oliveira

## Abstract

**Motivation:** With increasing initiatives to study biodiversity through high-quality reference genomes and the growing capacity of sequencing a wide range of organisms, there is a pressing need for an accessible, reproducible, and user-friendly tool that incorporates state-of-the-art methodologies for large genome assembly.

**Results:** We introduce Pipeasm, a Snakemake pipeline designed for assembling vertebrate genomes using HiFi PacBio, ONT, and HiC data. By setting a configuration file with input information and suggested parameters, Pipeasm was able to assemble multiple-sized diploid genomes ready for manual curation.

**Availability and Implementation:** Pipeasm requires an environment with Snakemake and Singularity and is available at https://github.com/itvgenomics/pipeasm.

## INTRODUCTION

Genome sequencing costs are steadily decreasing as the different technologies advance, providing the opportunity to investigate global biodiversity genomics in unprecedented detail. In contrast, the impact of the development of human society started the 6^th^ mass extinction, threatening most species on the planet (Ceballos, Ehrlich and Raven 2020). Thus, it is fundamental to develop tools to not only accelerate the generation, curation, and analyses of genomic information but also facilitate the use of such data in conservation efforts to appropriately comply with the connection through practical actions (Hogg *et al*. 2022; Hogg 2024).

Assessing the life on earth by genomics already presented its value for studying evolution (e.g., Feng *et al*. 2020; Zhang *et al*. 2020; Eizirik *et al*. 2023), conservation (e.g., Formenti *et al*. 2022b; Paez *et al*. 2022; Theissinger *et al*. 2023), and bioeconomy (e.g., Kreplak *et al*. 2019; Zhang *et al*. 2020; Peng *et al*. 2022). In this context, the creation of major initiatives to sequence and assemble the Earth’s biodiversity started to take form under umbrella projects like the Earth Biogenome Project (EBP) (Lewin *et al*. 2018; Gupta 2022). For assembling vertebrate genomes, the Vertebrate Genome Project (VGP) has been instrumental in defining the best approaches and metrics, unveiling species-specific characteristics that were previously unattainable (Rhie *et al*. 2021). Brazil’s initiative into this effort was born recently in two projects, the Genomics of the Brazilian Biodiversity (GBB) and the Genotropics (Genotropics 2023; Vilaça *et al*. 2024). Such endeavors have intrinsic difficulties, and the obstacles range from sample collection to computational infrastructure required to store and analyze the generated data, especially in the Global South.

One of the main challenges in generating reference genomes, as highlighted by the EBP, is the assembly and curation of massive genomic datasets on a large scale (Lewin *et al*. 2022). Therefore, it is crucial to develop and sustain modular pipelines to meet deadlines while ensuring adherence to quality standards and requirements (Lawniczak *et al*. 2022), making it accessible with diverse computational resources (Larivière *et al*. 2023). With ongoing advances in data generation and computational hardware, a range of bioinformatic tools have been developed to handle the various types of sequencing data (Ewels *et al*. 2020; Kuhl *et al*. 2020; Angelova *et al*. 2022; Larivière *et al*. 2023; Obinu *et al*. 2024). However, the multi-step assembly procedure can make the assembly process laborious, error-prone, and time-consuming.

Here, we present Pipeasm, an automated, customizable, and modular genome assembly pipeline designed in line with the best practices proposed in the context of the EBP and VGP to date. Requiring minimal manual intervention, the pipeline covers raw read trimming, overall genome statistics, contigging, and automated scaffolding, including genome assembly quality control after every major step, as well as benchmarking and logging. This workflow is built using Snakemake (Köster and Rahmann 2012), a powerful and user-friendly workflow management system, which makes Pipeasm adaptable, transparent, and easy to use in any Linux environment. By using Singularity (Kurtzer *et al*., 2017) containers, Pipeasm ensures that all dependencies are encapsulated, guaranteeing reproducibility and portability across different computing infrastructures. Configuration is streamlined and user-friendly, requiring only editing a single file to set up and customize the workflow, making it accessible to researchers with varying levels of computational expertise. Pipeasm is the main assembly strategy used in the GBB Project and is suggested for vertebrate diploid assemblies of any scale.

## METHODS

### Pipeline description

Pipeasm integrates a suite of tools to ensure high-quality data, from preprocessing steps to comprehensive reports and statistics. It is organized into six major steps: trimming and QC, k-mer profiling, assembly, assembly statistics, decontamination, and Hi-C mapping and scaffolding (Figure 1). The required input data are PacBio high-fidelity (HiFi-CCS) long reads, which enable solo assembly – an assembly performed using HiFi-only reads, delivering primary and alternate assemblies with partially phased contigs. Optionally, users can provide chromatin-contact (Hi-C) short reads to also perform phased assembly – which uses both HiFi and Hi-C to produce two haplotype-solved assemblies in ideal conditions (Cheng *et al*. 2021). When both data types are available, users can choose to perform only phased assembly for timing optimization or phased and solo assembly. Also, the user can provide Oxford Nanopore Long-Reads (ONT) to perform both solo and phased assemblies. Pipeasm offers a configuration file with parameters for all steps, already optimized with default values but easily customizable. The user must provide specific information within this file, including species name, sample ID, reads path, genetic code, taxonomy ID, and the BUSCO database to be used.

**Figure 1.**
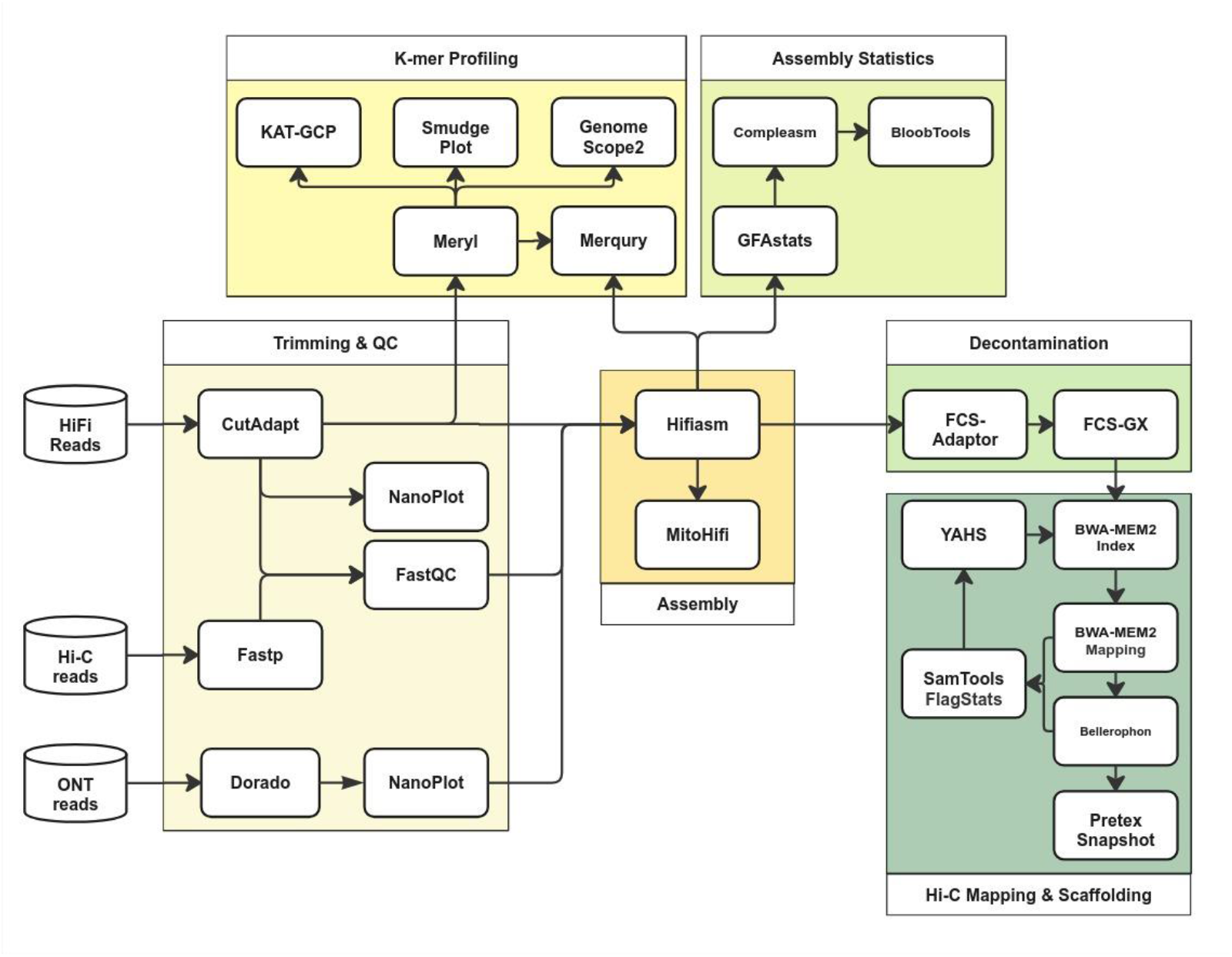
Schematic representation of the Pipeasm workflow, structured into six major tasks (large frames), each executed by the respective presented tools.

Pipeasm starts trimming PacBio HiFi-CCS long reads and Hi-C short reads using Cutadapt v4.4 (Martin 2011) and Fastp v0.23.4 (Chen *et al*. 2018), respectively, to guarantee high-quality input data for assembly. Dorado (https://github.com/nanoporetech/dorado) is used to trim all adapters and primers found in ONT reads. Quality control assessments are performed with FastQC v0.12.1 (Andrews 2023) for both long and short reads, while Nanoplot v1.41.6 (De Coster and Rademakers 2023) is used for long and ultra-long reads. Then, based on HiFi long reads, Pipeasm performs a set of analyses for detailed genomic statistics. For K-mer profiling, Meryl v1.3 (Rhie *et al*. 2020) is employed for K-mer counting and evaluation. GenomeScope v2.0 (Ranallo-Benavidez, Jaron and Schatz 2020) estimates genome size, heterozygosity, and repeat content, while SmudgePlot v0.3.0 (Ranallo-Benavidez, Jaron and Schatz 2020) identifies ploidy levels through K-mer distribution analysis. KAT-GCP v2.4.0 (Mapleson *et al*. 2017) enhances this analysis by providing graphical representations of K-mer spectra, aiding in identifying genomic features and contaminants.

The core assembly process is managed by Hifiasm v0.19.6 (Cheng *et al*. 2021), which produces haplotype-resolved genomes and is crucial for removing duplications between haplotypes and ensuring accurate, contiguous assemblies. Pipeasm also applies MitoHifi v3.2.2 (Uliano-Silva *et al*. 2023) to recover the mitogenome, ensuring accurate mitochondrial DNA assembly. GFAstats v1.3.6 (Formenti *et al*. 2022a) generates the fasta file from Hifiasm assembled graphs and detailed statistics on the assembled genome, including contig lengths, N50, and scaffold counts.

Post-assembly, Pipeasm integrates decontamination algorithms to eliminate remaining contaminants. FCS Adaptor v0.5.0 removes adaptors, while FCS-GX v0.5.0 targets other contaminants (Astashyn *et al*. 2024). The assembly quality control is evaluated through gene-space completeness, assembled K-mers compared to the read’s K-mer distribution, and overall assembly metrics. Compleasm v0.2.2 assesses genome completeness by comparing the assembled genome against BUSCO databases to evaluate the presence of conserved single-copy orthologs (Manni *et al*. 2021; Huang and Li 2023). Merqury v1.3 (Rhie *et al*. 2020) performs the assembled K-mer counts and compares them to the Meryl database created in previous steps. For intuitive visualization, a snailplot, obtained with BlobToolkit v4.3.5 (Challis *et al*. 2020), provides a comprehensive overview of the assembled genome.

To elucidate chromosomal interactions, Hi-C reads are mapped onto phased genome assemblies. Genome indexing and mapping are performed using BWA-MEM2 v2.2.1 (Vasimuddin *et al*. 2019) for both haplotypes from decontaminated assemblies. Subsequently, forward and reverse Hi-C reads are aligned separately to each haplotype and sorted with SAMtools v1.19 (Danecek *et al*. 2021). Low-quality alignments and duplicated paired-end reads are filtered and then merged using Bellerophon v1.0 (https://github.com/davebx/bellerophon), consolidating interaction information into single BAM files for each haplotype.

Quality assessment of the above-mentioned mapping and its merging process is conducted using SAMtools flagstat and the script get_stats.pl (https://github.com/ArimaGenomics/mapping_pipeline), providing essential metrics such as alignment rates, duplication levels, and distance between contact read pairs. Afterwards, Pretext tools (Map v0.1.9 and Snapshot v0.0.4) (https://github.com/sanger-tol/PretextMap, https://github.com/sanger-tol/PretextSnapshot) convert BAM files into visual representations of Hi-C contact maps. Finally, the automatic scaffolding step, which was performed using YAHS v1.0 (Zhou, McCarthy and Durbin 2023), links contigs into longer scaffolds, enhancing the assembly. The entire mapping step is repeated against the scaffolded genome to generate a final contact map. Then, the last steps of quality check are performed, over the scaffolded genome with GFAstats, Compleasm and Merqury.

Pipeasm also compiles multiple metrics and quality assessments from all stages of the genomic assembly process into comprehensive output files. These files encapsulate essential information, such as read quality statistics, genome size estimations, contamination analyses, and assembly quality metrics. The output includes summaries of genome completeness and contiguity, highlighting key indicators such as N50 values, total base pairs, and the presence of conserved genes. Additionally, quality values (QV) and BUSCO scores provide insights into the accuracy and completeness of the assembled genome

The steps mentioned above are conducted and managed using the Snakemake workflow engine, following its best practices and a standardized repository structure to ensure reproducibility, adaptability, and transparency (Mölder *et al*. 2021). This Python-based approach offers easy readability and automatically scales Pipeasm tasks for parallelization, logging, and benchmarking. We are also providing a bash script for easy usage and allowing users to split the run of each step of the pipeline.

### Computational environment and running test

The test environment for running Pipeasm used an Intel^(R)^ Xeon^(R)^ Gold 6252 CPU @ 2.10GHz, leveraging 64 threads overall, with 32 threads allocated for each rule within Snakemake. Performance metrics and resource utilization were systematically monitored and extracted using Snakemake’s built-in benchmark tool, ensuring accurate assessment and optimization of computational efficiency throughout the process. It is important to note that the computation time reported is the sum of all CPU time used by all rules rather than the straight running time, providing a comprehensive measure of the workflow’s computational demand.

We employed genomic data from 4 different species: (A) *Gallus gallus* (bGalGal), a chicken from the Phasianidae family, with a diploid genome size of 1.1 Gbp and 39 chromosomes + WZ (NCBI Genome Accession GCA_027557775.1). Reads were downloaded from the bGalGal5 individual, with 45.88 Gbp PacBio HiFi reads and 147.73 Gbp Arima Hi-C reads. (B) *Taeniopygia guttata* (bTaeGut), a bird from the Estrildidae family, with a diploid genome size of 1.1 Gbp and 37 chromosomes + WZ (NCBI Genome Accession GCA_003957565.4). Reads downloaded from the bTaeGut2 individual have 39.52 Gbp PacBio HiFi reads and 99.04 Gbp Arima Hi-C reads. (C) *Elephas maximus* (mEleMax), a mammal from the Elephantidae family, with a diploid genome size of 3.4 Gbp and 27 chromosomes + XY (NCBI Genome Accession GCA_024166365.1). Reads downloaded from the mEleMax1 individual have 137.82 Gbp PacBio HiFi reads and 277.42 Gbp Arima Hi-C reads. (D) *Pseudophryne corroboree* (aPseCor), a frog from the Myobatrachidae family, with a diploid genome size of 8.9 Gbp and 12 chromosomes (NCBI Genome Accession GCA_028390025.1). Reads downloaded from the aPseCor3 individual have 230.31 Gbp PacBio HiFi reads and 381.02 Gbp Arima Hi-C reads. All above mentioned reads were downloaded from Genome Ark (https://www.genomeark.org).

## RESULTS AND DISCUSSION

Pipeasm successfully assembled the data from bGalGal, bTaeGut, mEleMax and aPseCor (Table 1), reflecting expected biological patterns. Together with k-mer profile analyses (Figure 2), the assemblies presented the expected ploidy and range of genome size, as estimated by GoaT, previous assemblies and related species literature (Formenti *et al*. 2021; Rhie *et al*. 2021; Challis *et al*. 2023). In general, haplotypes from the same assembly produced similar statistics, with minor differences in repeat regions and the presence of sex chromosomes, as observed on the contact maps and summary results (Figure 3, Table 1). When examining the genome characteristics of the individuals, bTaeGut showed the highest heterozygosity (1.34%) and error estimates (0.267), while aPseCor had the largest genome size (8.83 Gb and 8.97 Gb for haplotypes one and two, respectively) and the most repetitive genome (65.5%). Notably, only in mEleMax did the scaffolding step increase the number of scaffolds (from 571 and 852 in the phased assembly to 642 and 954 post-scaffolding for the primary and alternative assemblies, respectively), but it also improved overall contiguity. When comparing the Pipeasm assembly to their respective first-version assemblies, we observed similar or improved statistics (Figure 4). Given that Pipeasm follows the latest best practices from the same consortium that assembled these genomes, differences in algorithms, software versions, and different purging steps may explain the observed improvements.

**Table 1:** Statistics for genome assembly of aPseCor, bGalGal, bTaeGut, and mEleMax (Hifiasm phased mode followed by YAHS) for both Primary (haplotype 1) and Alternative (haplotype 2). The main genome assembly metrics were calculated with gfstats, while completedness was estimated by Compleasm with OrthoDB version 10 verifying: tetrapoda orthologs for aPseCor (5310 genes), birds orthologs for bGalGal (8337 genes), passeriformes orthologs for bTaeGut (10844 genes), and eutheria orthologs for mEleMax (11636 genes).

| Genome Stats | aPseCor |  | bGalGal |  | bTaeGut |  | mEleMax |  |
| --- | --- | --- | --- | --- | --- | --- | --- | --- |
|  | Hap1 | Hap2 | Hap1 | Hap2 | Hap1 | Hap2 | Hap1 | Hap2 |
| **Scaffolds** | 2,535 | 2,454 | 565 | 270 | 434 | 210 | 642 | 954 |
| **Size Length** | 8,83 Gb | 8,97 Gb | 1,13 Gb | 981 Mb | 1,17 Gb | 995 Mb | 3,26 Gb | 3,63 Gb |
| **Largest Scaffold** | 46,04 Mb | 49,01 Mb | 33,08 Mb | 36,57 Mb | 30,72 Mb | 25,22 Mb | 175,39 Mb | 203,88 Mb |
| **Scaffold N50** | 918,47 Mb | 821,71 Mb | 86,73 Mb | 91,13 Mb | 72,12 Mb | 71,94 Mb | 116,12 Mb | 116,40 Mb |
| **Scaffold L50** | 4 | 4 | 5 | 4 | 6 | 5 | 10 | 11 |
| **GC (%)** | 45.67 | 45.6 | 42.6 | 42.26 | 42.92 | 42.37 | 41.32 | 41.55 |
| **Gaps** | 2258 | 2263 | 438 | 291 | 442 | 266 | 52 | 63 |
| **Single copy (%)** | 95.35 | 95.50 | 99.57 | 95.36 | 99.63 | 99.91 | 97.11 | 99.43 |
| **Duplicated (%)** | 0.49 | 0.55 | 0.17 | 0.14 | 0.12 | 0.06 | 0.31 | 0.31 |
| **Fragmented (%)** | 0.83 | 0.70 | 0.02 | 0.11 | 0.06 | 0.06 | 0.08 | 0.03 |
| **Missing (%)** | 3.33 | 3.26 | 0.24 | 4.39 | 0.18 | 0.98 | 2.51 | 0.24 |

**Figure 2.**
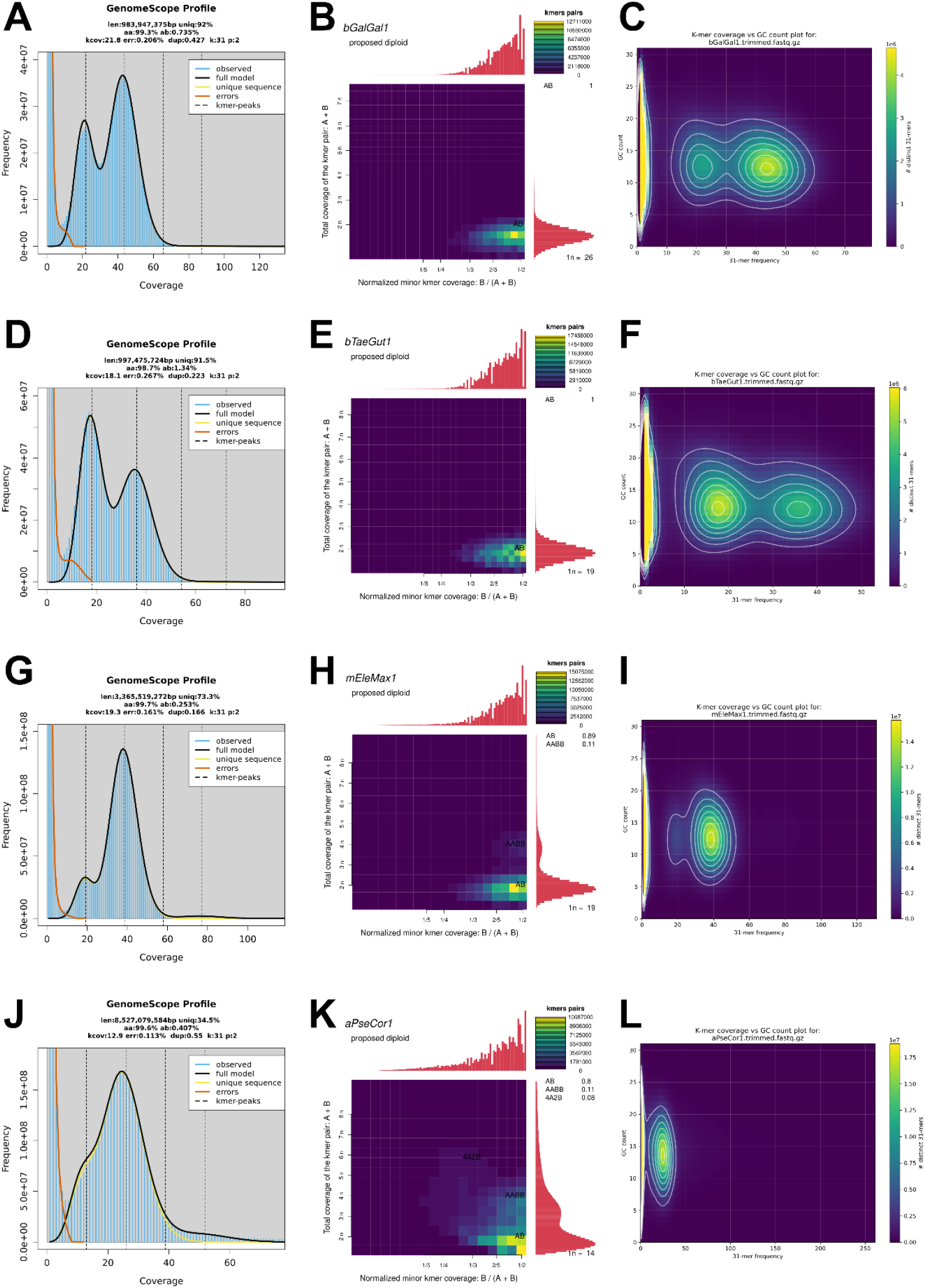
Summary metrics of species genomes based on k-mer distribution. Panels A-C stand for bGalGal, D-F for bTaeGut, G-I for mEleMax and J-L for aPseCor. Left panels are GenomeScope2 results, showing predicted genome length, duplication rate, heterozygosity. Middle panels are SmudgePlot results, showing estimated ploidy and k-mer depth. The right panels are KAT-GCP results showing k-mer frequency per GC count. All estimates presented used k=31. (A) bGalGal GenomeScope2 analysis showing predicted genome length of 983,947 MB, 92% uniq, 99.3% aa, 0.735% ab, 21.8 kcov, 0.206% err, 0.427 dup, kmer 31 and ploidy of 2. (B) SmudgePlot of bGalGal showing 1n = 26 and proposed diploid. (D) bTaeGut GenomeScope2 analysis showing predicted genome lenght of 997,475 MB, 91.5% uniq, 98.7% aa, 1.34% ab, 18.1 kcov, 0.267% err, 0.223 dup, kmer 31 and ploidy of 2. (E) SmudgePlot of bTaeGut showing 1n = 19 and proposed diploid. (G) mEleMax GenomeScope2 analysis showing predicted genome lenght of 3,365,519 MB, 73.3% uniq, 99.7% aa, 0.253% ab, 19.3 kcov, 0.161% err, 0.166 dup, kmer 31 and ploidy of 2. (H) SmudgePlot of mEleMax showing 1n = 19 and proposed diploid. (J) aPseCor GenomeScope2 analysis showing predicted genome lenght of 8,527,079 MB, 34.5% uniq, 99.6% aa, 0.407% ab, 12.9 kcov, 0.113% err, 0.55 dup, kmer 31 and ploidy of 2. (K) SmudgePlot of aPseCor showing 1n = 14 and proposed diploid. C, F, I and L are for KAT-GCP plot showing 31 k-kmer frequency per GC count and distinct 31-mers.

**Figure 3.**
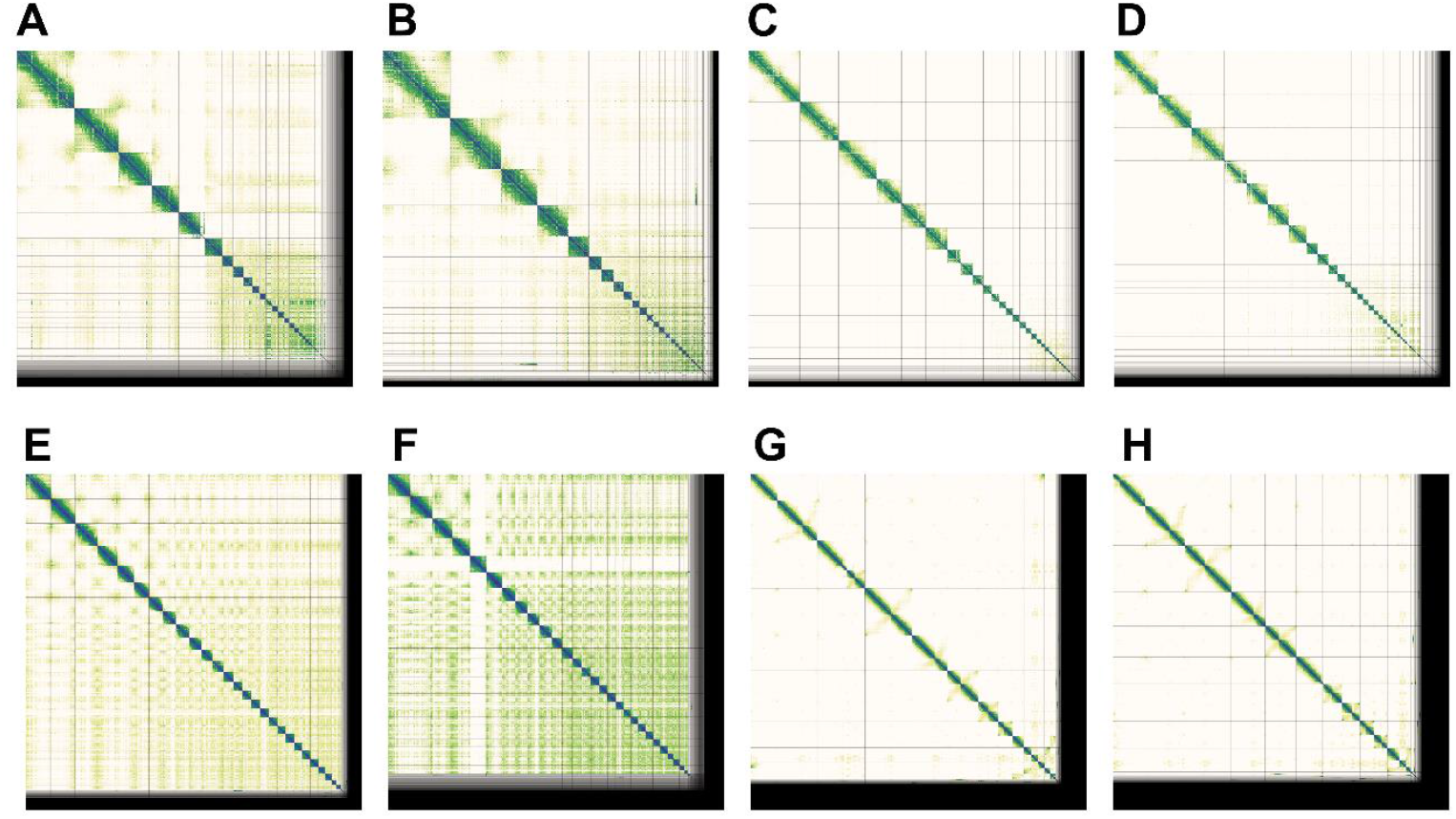
Summary of Hi-C mapping snapshots. Pretext snapshots of both haplotype 1 (top) and haplotype 2 (bottom) of all assemblies are shown. Panel orders A-D and E-H stand for bGalGal, bTaeGut, mEleMax and aPseCor, respectively.

**Figure 4.**
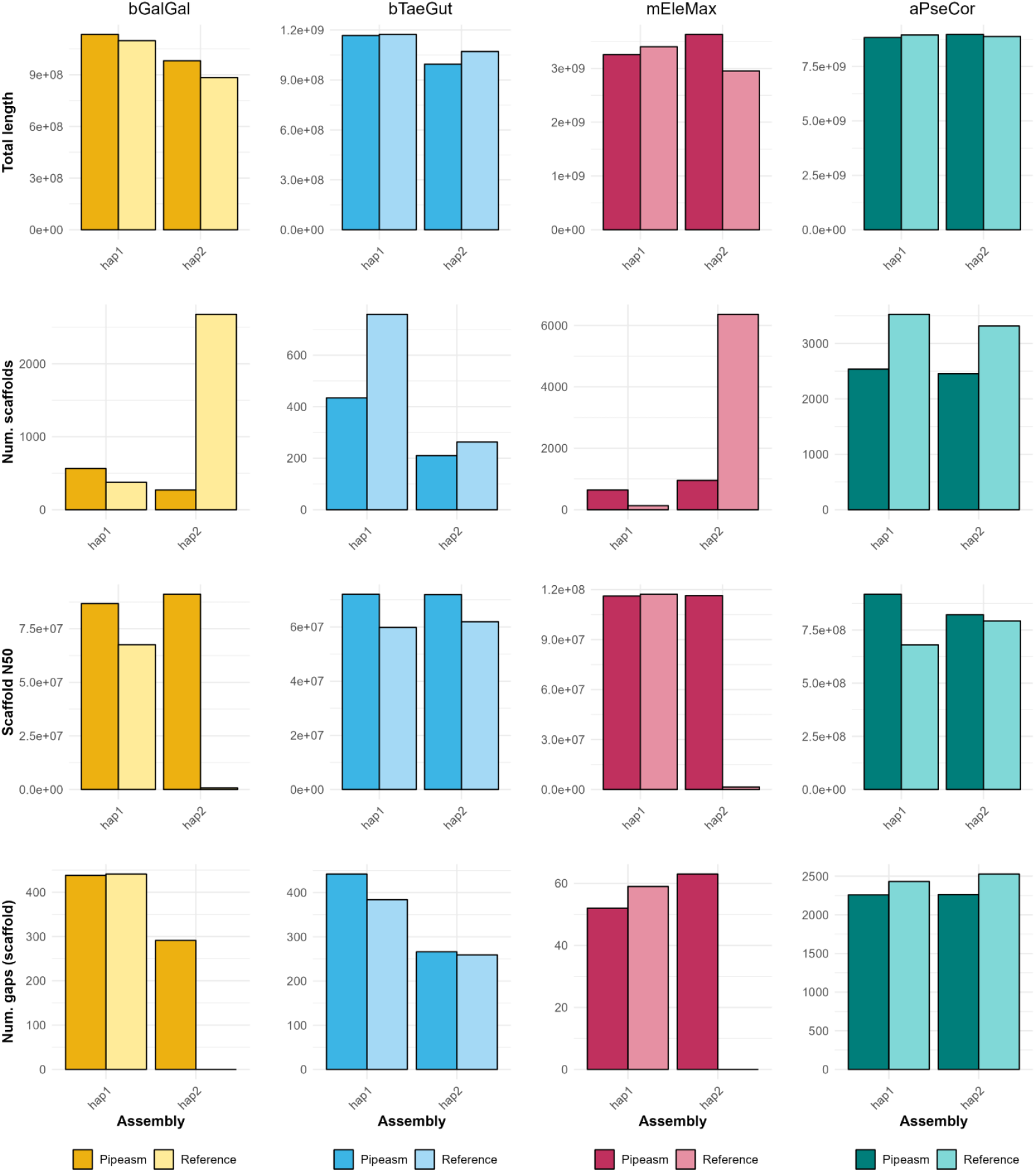
Summary metrics of scaffolding assembly. Key metrics for both the reference and the resulting Pipeasm scaffolding assembly process are presented. Each row represents a metric, and each column corresponds to a species assembly. Note that the plots’ Y axes are not within the same scale. The first and second haplotypes from different pipelines are not necessarily completely equivalent.

Benchmarking results for the above assemblies provided insights into resource utilization and computational efficiency across various steps of Pipeasm (Figure 5). The entire process for a solo and phased assembly, including scaffolding and Hi-C maps, took approximately 23 hours for bTaeGut. Running time is mainly affected by genome length, although it is important to note that it also depends on the computational infrastructure. All our assemblies were conducted in a High-Performance Computing (HPC) environment, and the running times may vary based on factors such as disk input/output operations, CPU clock speed, and memory frequency. Still, the relationship between CPU time and RAM usage will likely remain consistent across different setups. Overall, the observed running time is primarily due to the optimization provided by Snakemake in managing task execution, since Pipeasm: (i) allocates fewer cores to simple tasks that do not serve as prerequisites for others; and (ii) automatically initiates new tasks that can be run concurrently without waiting for previous tasks to be completed or requiring user intervention to start subsequent tasks.

**Figure 5.**
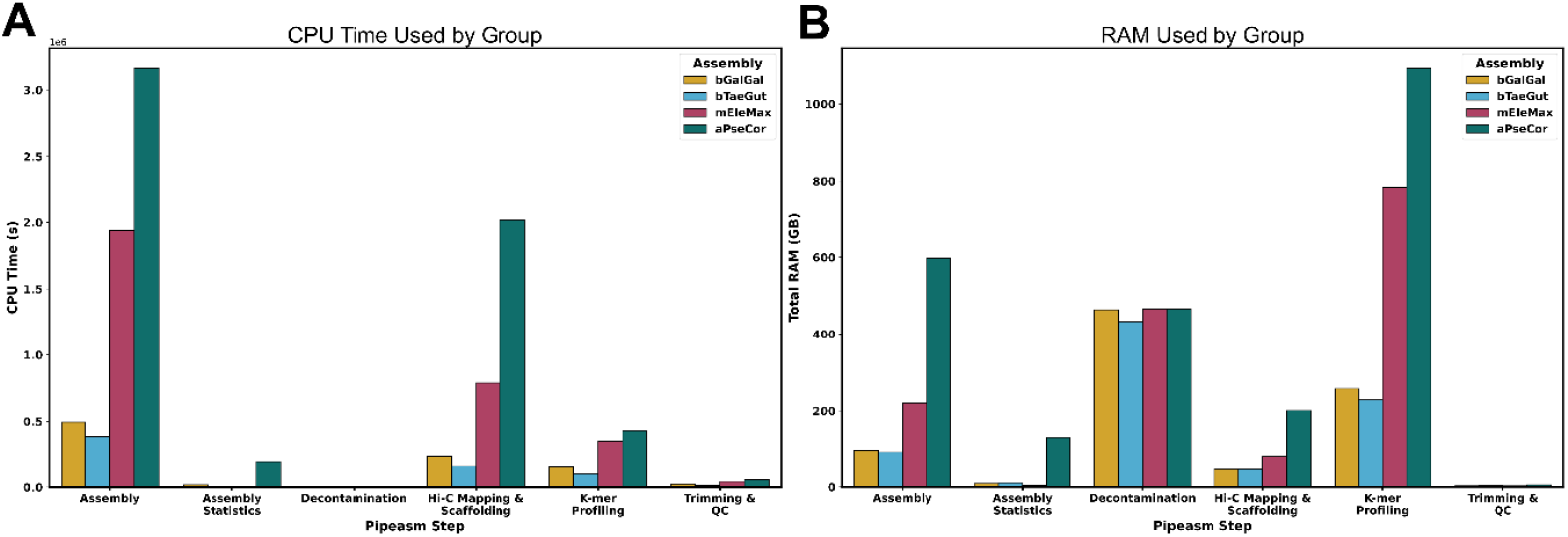
Pipeasm Performance Metrics: CPU and RAM Usage Analysis. Detailed analysis of CPU and RAM usage across various stages of Pipeasm’s combined steps. (A) CPU time consumption across different pipeline steps, including Assembly, Assembly Statistics, Decontamination, Hi-C Mapping & Scaffolding, K-mer Profiling, and Trimming & QC. The results are segmented into four different assembly groups: bGalGal, bTaeGut, mEleMax, and aPseCor. The mEleMax group exhibits significantly higher CPU time usage, particularly in the Assembly and Hi-C Mapping & Scaffolding steps. (B) illustrates the total RAM usage for the same pipeline steps and assembly groups. The K-mer Profiling step shows the highest RAM consumption, especially for the mEleMax group, indicating a resource-intensive process.

The assembly process was time and memory-intensive, with the mEleMax and aPseCor requiring significant CPU time and RAM (Figure 5), particularly for aPseCor, the largest genome size (Table 1). The decontamination step also demanded considerable memory for all species once it used the same FCS-GX database for sequence alignment. The most resource-heavy stage was Hi-C mapping and scaffolding, which required extensive CPU time but had moderate memory use. K-mer profiling had the highest memory consumption but took less CPU time, and adjusting settings in Meryl could help reduce RAM usage. The trimming and quality control steps, though requiring less memory, still took a notable amount of processing time, especially for aPseCor. Since Pipeasm requires HiFi reads for assembly, it meets the minimum requirements for achieving high-quality vertebrate genome assemblies. Including Hi-C and/or ONT data and performing phased assembly further elevates the pipeline to higher standards (Rhie *et al*. 2021). Although the scaffolding step is the most time-consuming (Larivière *et al*. 2023), it is essential for producing genomic information that is ready for manual curation. Moreover, during the execution of the entire pipeline, users can monitor task completion, review output files, and check the analysis logs, ensuring a high degree of transparency (Mölder *et al*. 2021). The evaluation and adjustment of parameters can be achieved without requiring the entire pipeline to be completed before re-running any steps. Tasks that have already been performed are automatically recognized by Snakemake, ensuring that only the necessary steps are re-executed. Indeed, the recent implementation of numerous pipelines for high-throughput sequencing data using Snakemake (e.g. Mölder *et al*. 2021; Mohsen *et al*. 2022; Fallon *et al*. 2023; Gregoricchio and Zwart 2023; Neuenschwander *et al*. 2023; Weber, Cosenza and Korbel 2023), already underscores its significant value.

When compared to recently published assembly pipelines (e.g. Kuhl *et al*. 2020; Angelova *et al*. 2022; Larivière *et al*. 2023), Pipeasm presents several differences and advantages. SnakeCube, for instance, does not utilize HiFi or Hi-C data and employs different tools and approaches for assembly, including a polishing step that is less common in current methodologies (Angelova *et al*. 2022). CSA also does not use HiFi data and requires an additional re-assembly step. It lacks genome assembly evaluation during the process, and being a Perl script, it may be less transparent, flexible, and readable for some users. While CSA offers good performance in terms of time consumption and the ability to input data from additional species, it employs a methodology that is not widely adopted today (Kuhl *et al*. 2020).

The Galaxy pipeline (Larivière *et al*. 2023) adheres to VGP standards and follows a similar concept to Pipeasm. However, Pipeasm’s implementation using Snakemake and Singularity offers greater flexibility, adaptability, readability, modularity, and transparency, particularly for users with limited coding skills. Furthermore, the Galaxy pipeline, whether used online or locally, can be very limiting depending on the region, data, and resources available to the researcher. In contrast, Pipeasm provides a more versatile and accessible solution for various genomic assembly projects.

The Sanger nf-core genomeassembly tool (Ewels *et al*. 2020; Krasheninnikova *et al*. 2023) developed with NextFlow, and Colora (Obinu *et al*. 2024), another pipeline in Snakemake, are also available for genome assembly. However, Pipeasm remains the only one that delivers a genome ready for manual curation, with contact maps generated and a pretext file for scaffold editing. Unlike the others, Pipeasm also provides various k-mer analyses, statistics, and publication-ready figures and reports. Our pipeline stands out for being the most user-friendly, as it does not require manual downloading of any database if you choose not to run the genome decontamination step.

## CONCLUSIONS

Pipeasm has proven to meet all expectations regarding the optimization of analyses and reduction of running time – when compared to running each step separately – while maintaining high assembly quality. The vertebrate species used in this study serve as surrogates to the ones to be included in the GBB project, demonstrating that Pipeasm delivers a high-quality genome assembly ready for curation within approximately one to four straight days, depending on the specific genome structure and available computational resources. The tool also provides a straightforward approach to obtain the most relevant results for evaluating the assembly.

Furthermore, the use of Snakemake endows Pipeasm with valuable readability, modularity, adaptability, and transparency. Users can easily determine which commands will be executed and, with basic coding skills, can add new features according to their needs. Additionally, by utilizing Singularity to containerize the required software, Pipeasm simplifies the provision of all necessary bioinformatics dependencies without installation and facilitates the seamless updating of tools to newer versions or even their replacement with alternative tools as needed.

Pipeasm presents limitations that are intrinsic to the tools used, such as the need for enough computational resources and its restricted applicability to certain organisms. Future iterations of Pipeasm will include verifying and adapting tools to apply the pipeline to other taxa (e.g., plants) and providing basic modules with key analyses recommended by the Earth BioGenome Project (EBP) (Lawniczak *et al*. 2022). Additionally, there is an urgent need for an annotation pipeline, given the challenges observed in reference genome initiative projects (Lewin *et al*. 2022). Currently, Pipeasm significantly facilitates the assembly of vertebrate reference genomes. Its simplicity of use and focus on providing the most useful results directly contribute to the popularization of genomic information. This is a crucial step for large-scale projects focusing on conservation, biodiversity studies, and applications in the bioeconomy.

